# Automated Segmentation of Prostatic Gold Fiducial Markers for MR-Only Radiotherapy Planning Using Multi-Model Consensus Deep Learning

**DOI:** 10.64898/2026.06.18.733061

**Authors:** Ashley W. Stewart, Jonathan Goodwin, Matthew Richardson, Simon D. Robinson, Kieran O’Brien, Jin Jin, Markus Barth, Steffen Bollmann

## Abstract

**Purpose:** To develop and evaluate a multi-model consensus deep learning approach for automated gold fiducial marker (FM) segmentation in T1-weighted prostate MRI.

**Materials and Methods:** In this retrospective study, T1-weighted MRI and CT-derived reference standard segmentations were collected from 127 prostate cancer patients (all male; mean age, 70 years ± 7 [standard deviation]; age range, 50–88 years; collected between October 2020 and January 2026) who each had three implanted gold FMs. A 3D U-Net was trained on 93 subjects using four random seeds to produce an ensemble. At inference, marker-class probability maps were averaged across models and the top three connected components selected. Performance was evaluated on 34 temporally held-out subjects (9 tuning, 25 test) using marker-level sensitivity and precision with exact (Clopper-Pearson) 95% confidence intervals (CIs). A model count ablation study was performed. The pipeline was deployed for on-scanner processing on Siemens MRI systems via the OpenRecon framework and as a browser-based application using WebAssembly, executing entirely client-side.

**Results:** The four-model consensus achieved 96% (70 of 73) sensitivity and 95% (70 of 74) precision on 25 test subjects, with 29 of 34 (85%) subjects achieving perfect marker detection. Single models had a mean sensitivity of 84% (SD, 9%), improving to 96% with four-model consensus (SD, *<*1%).

**Conclusion:** Multi-model consensus deep learning substantially improved FM segmentation reliability over individual models, achieving high sensitivity and precision using only routinely acquired T1-weighted MRI.

## 1 Introduction

Prostate cancer is the second most common cancer in men, with over 1.2 million new cases and 350,000 deaths annually.^1^ For localized disease, intensity-modulated radiotherapy targets tumor volumes while minimizing dose to surrounding tissues, but geometric misses can occur due to inter- and intrafractional patient motion.^2,3^ Gold fiducial markers (FMs) implanted into the prostate gland under ultrasound guidance mitigate these errors by enabling marker-based image-guided radiotherapy.^4,5^ In a CT-MRI planning workflow, MRI provides superior soft-tissue contrast for target delineation while CT provides attenuation information for dosimetry and confident FM visualization.

Interest in MR-only radiotherapy planning has grown considerably,^6–8^ as it offers reduced uncertainty by eliminating CT-MR registration, a streamlined patient pathway, reduced ionizing radiation exposure, and alleviates demand on CT resources. Synthetic CT derived from MRI addresses the dosimetry requirement.^9–12^ However, accurate FM detection in MRI remains a key challenge because FMs appear as signal voids that are difficult to differentiate from calcifications and other void sources.

Automated FM detection techniques for MR-only radiotherapy have progressed from template-matching methods achieving 63–88% accuracy,^13^ through simulated complex-valued templates reaching 96%,^14^ to multiparametric approaches achieving 84–94% true-positive rates.^15,16^ Deep learning advanced performance further, with Gustafsson et al.^17^ demonstrating 97.4% detection sensitivity across 39 patients. Quantitative susceptibility mapping (QSM) has also been explored for FM differentiation from calcifications,^18^ but requires specialized sequences not routinely available. In this work, we present a multi-model consensus deep learning approach for automated FM segmentation using only T1-weighted MRI. The contributions are (Fig 1): (1) training on a large cohort of 127 patients; (2) a consensus inference strategy averaging probability maps across independently trained models; (3) evaluation on 34 held-out subjects; and (4) deployment for on-scanner processing via the Siemens OpenRecon framework and as a browser-based application using WebAssembly.

**Figure 1.**
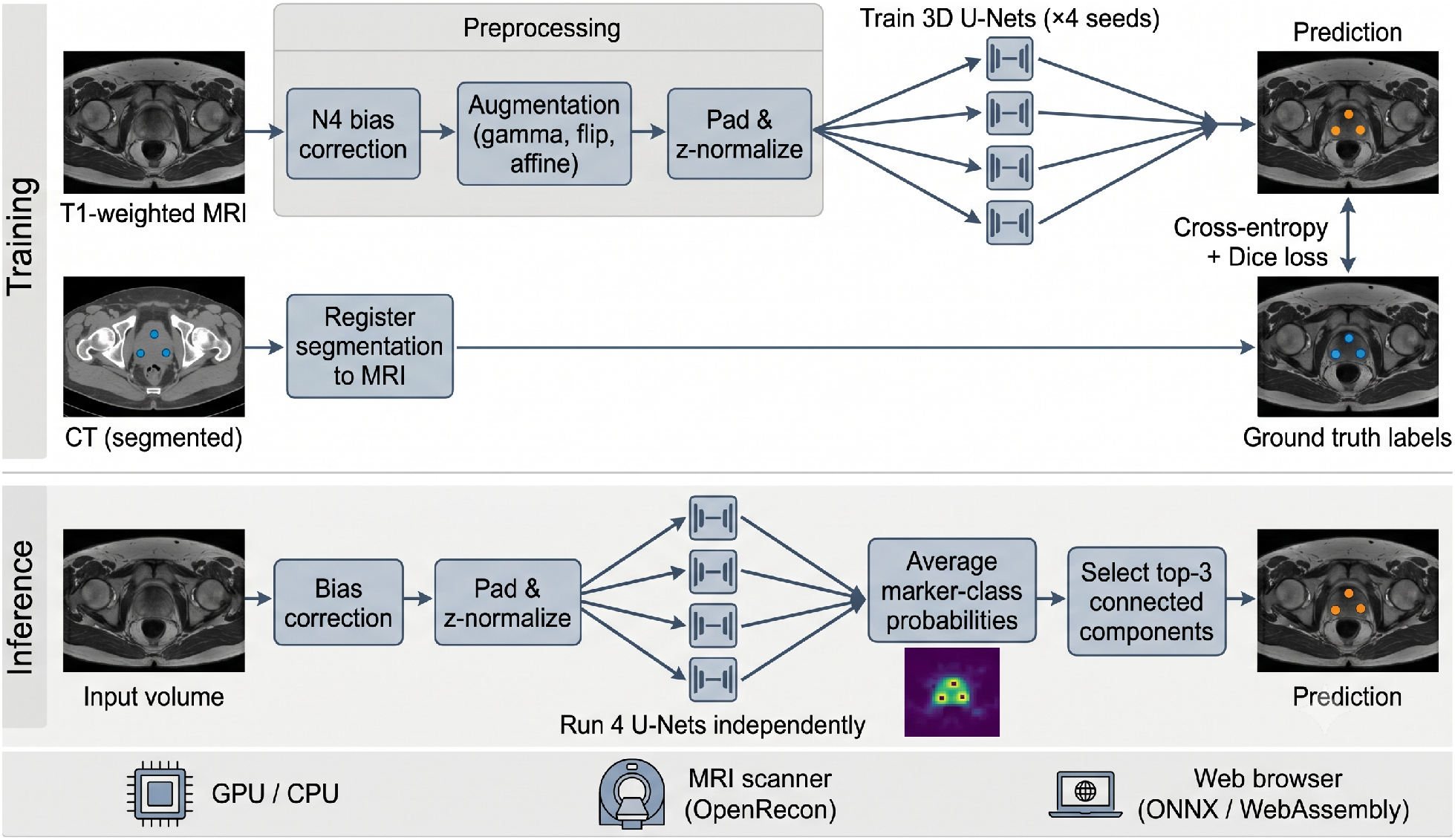
Overview of the training and inference pipelines. Top: T1-weighted MRI volumes are preprocessed and used to train four 3D U-Net models with different random seeds, supervised by CT-derived reference standard segmentations registered to MRI space. Bottom: at inference, the four models independently process the input volume, their marker-class probability maps are averaged, and the top three connected components are selected. The same pipeline runs on GPU/CPU, on the MRI scanner via OpenRecon, and in the web browser via ONNX/WebAssembly.

## 2 Materials and Methods

### 2.1 Ethics and Data Collection

This retrospective study was approved by the relevant regional ethics committee (approval number available on request), with site-specific institutional authorization. The requirement for written informed consent was waived due to the retrospective nature of the study. All imaging data were de-identified prior to analysis in accordance with institutional data governance policies.

MRI and CT data were collected retrospectively from 127 prostate cancer patients (all male; mean age, 70 years ± 7; age range, 50–88 years), each with three implanted gold FMs (Riverpoint Medical), cylindrical and 1 × 3 mm. Data were acquired between October 2020 and January 2026.

Age was available for 126 of 127 subjects with recoverable dates of birth.

### 2.2 Image Acquisition

CT data were acquired on a Siemens SOMATOM Confidence scanner (Siemens Healthineers) with resolution of 0.98 × 0.98 × 2 mm, matrix size of 512 × 512 × 188, and tube voltage of 120 kV.

T1-weighted MRI data were acquired on a 3T MAGNETOM Skyra scanner (Siemens Healthineers) with a 32-channel spine coil and 18-channel body coil. A T1-weighted gradient-echo sequence was used with repetition time msec/echo time msec, 600/6.66; flip angle, 80°; field of view, 220 × 220 mm; matrix, 320 × 320; in-plane resolution, 0.69 × 0.69 mm; section thickness, 2.0 mm; 40 sections; and parallel imaging acceleration factor of 2.

### 2.3 Reference Standard

FM and clinical target volume (CTV) segmentations were delineated on registered CT images by radiation oncologists as part of routine clinical treatment planning and transformed to MRI coordinates using Digital Imaging and Communications in Medicine (DICOM) registration objects via SimpleITK. Individual marker regions of interest and the CTV were rasterized to create a three-class segmentation label: background (class 0), gold seed markers (class 1), and prostate (class 2). All T1-weighted volumes were preprocessed with N4 bias field correction.^19^

### 2.4 Dataset Partitions

The 127 subjects were divided into three nonoverlapping sets: 93 for training, 9 for tuning (used for early stopping), and 25 for testing. The tuning and test subjects were drawn from a temporally separate clinical cohort (August 2025 to January 2026), while training data spanned earlier acquisitions (October 2020 to 2025), providing a temporal held-out evaluation. No test data were used during model development or hyperparameter selection.

### 2.5 Model Architecture

A 3D U-Net was implemented in PyTorch 2.1.0 (Meta AI) with CUDA 12.1. The encoder comprised five levels with channel counts of 16, 32, 64, 128, and 256, and a bottleneck of 512 channels. The decoder mirrored the encoder with skip connections. The network accepted single-channel input (N4-corrected T1-weighted MRI) and produced three-class output. Input volumes were padded to the nearest multiple of 32 in each dimension, and intensities were standardized using z-score normalization.

### 2.6 Training

The loss function combined weighted cross-entropy (class weights of 1, 5, and 1 for background, markers, and prostate, respectively) and Dice loss with equal weighting. The Adam optimizer was used with an initial learning rate of 0.003 and cosine annealing schedule. Training ran for up to 1300 epochs with early stopping triggered after 500 epochs without improvement in tuning loss.

Data augmentation included random intensity shifts (gamma correction), random flipping along all axes, random affine transformations (rotations up to 90°), and z-score normalization via TorchIO 0.18.91.^20^ The batch size was 6. Four models were trained on the full training set using different random seeds (42, 123, 456, 789), with each model evaluated against the same 9 tuning subjects. Training was performed on NVIDIA A100 GPUs, with each model converging in 663–868 epochs (approximately 7–10 hours).

### 2.7 Consensus Inference

At inference, each model independently produced class probability maps via softmax. The marker-class probability maps were averaged across all four models to produce a consensus map. Connected components were identified in the thresholded consensus map (threshold = 0.1), and the top three components ranked by mean probability were selected as the final predictions. This top-3 selection reflects the clinical prior that each patient has exactly three implanted FMs.

### 2.8 Evaluation Metrics

Performance was assessed at the marker level using connected-component matching. Both predicted and reference standard segmentations were dilated by one voxel (3 × 3 × 3 structuring element) before component labeling to account for minor spatial offsets. A target marker was counted as detected if any predicted component overlapped it after dilation. Sensitivity was defined as detected markers divided by total markers, and precision as true detections divided by total predictions. A subject had perfect detection if all markers were found with no false positives.

### 2.9 Model Count Ablation

To quantify the benefit of multi-model consensus, performance was evaluated using one, two, three, and four models. For subsets of fewer than four models, all possible combinations were evaluated and mean ± standard deviation (SD) reported.

### 2.10 Deployment

The segmentation pipeline was deployed for on-scanner processing on Siemens MRI systems using the OpenRecon framework,^21^ which enables containerized reconstruction algorithms to run on-scanner during the clinical workflow. The pipeline was packaged as a container using the NeuroDesk project’s neurocontainers infrastructure,^22,23^ which defines reproducible container recipes as declarative specifications. The container communicates with the scanner using the standardized MRI raw data protocol MRD,^24^ performing N4 bias field correction, multi-model consensus segmentation, morphological post-processing, and overlay of detected marker contours onto the original images for immediate visualization on the scanner console. The container runs with and without GPU acceleration, ensuring compatibility with various scanner reconstruction hardware. The build and deployment pipeline is automated via continuous integration: recipe updates trigger automatic rebuilding and distribution to scanner-deployable archives.

Additionally, the consensus pipeline was implemented as a browser-based application (https://seedseg.neurodesk.org) executing entirely client-side using Open Neural Network Exchange (ONNX) Runtime Web^25^ with a WebAssembly backend, preserving patient data privacy by design. Trained model weights were converted to ONNX format (approximately 87 MB each) and cached in the browser’s IndexedDB storage after first download. N4 correction was implemented using a WebAssembly build of MriResearchTools.jl,^26^ and DICOM conversion via dcm2niix.^27^ The application requires no software installation and accepts both NIfTI and DICOM inputs; an interactive viewer based on NiiVue^28^ allows inspection of input volumes and segmentation results. When DICOM inputs are provided, the application can validate the imaging protocol against expected acquisition parameters using dicompare,^29^ allowing users to assess whether sequence differences may affect segmentation. No data are uploaded to external servers.

### 2.11 Statistical Analysis

Sensitivity and precision were calculated as proportions with numerators and denominators reported. Exact 95% confidence intervals (CIs) were computed using the Clopper-Pearson method. A *P* value less than .05 indicated a statistically significant difference. Age differences between partitions were compared using the Welch *t* test. For the model count ablation, all possible model combinations were evaluated for each subset size (*C*(4, *k*) combinations for *k* models), and mean ± SD across combinations was reported. Per-subject sensitivity under four-model consensus was compared to the mean per-subject sensitivity across the four single-model inferences using a Wilcoxon signed-rank test. Bootstrap 95% CIs (10,000 resamples) were computed for sensitivity at each model count. Analyses were performed using Python 3.9 (Python Software Foundation) with SciPy 1.13.1 (scipy.org). Preprocessing used SimpleITK 2.4.1, nibabel 5.3.2, and pydicom 2.4.4.

### 2.12 Data and Code Availability

All training, evaluation, and analysis code is available on GitHub: https://github.com/astewartau/seedseg. The browser-based inference application is available at https://seedseg.neurodesk.org. The containerized deployment pipeline is available through the NeuroDesk project: container recipe at https://github.com/NeuroDesk/neurocontainers and OpenRecon packaging at https://github.com/NeuroDesk/openrecon. Trained model weights are hosted on the Open Science Framework (https://osf.io/f7hjv/). The imaging data are not publicly available due to patient privacy restrictions but may be available upon reasonable request to the corresponding author.

## 3 Results

### 3.1 Training Convergence

All four models converged to similar losses despite different random initializations (Fig 2). Early stopping was triggered between epochs 663 and 868. Loss curves showed stable convergence without overfitting.

**Figure 2.**
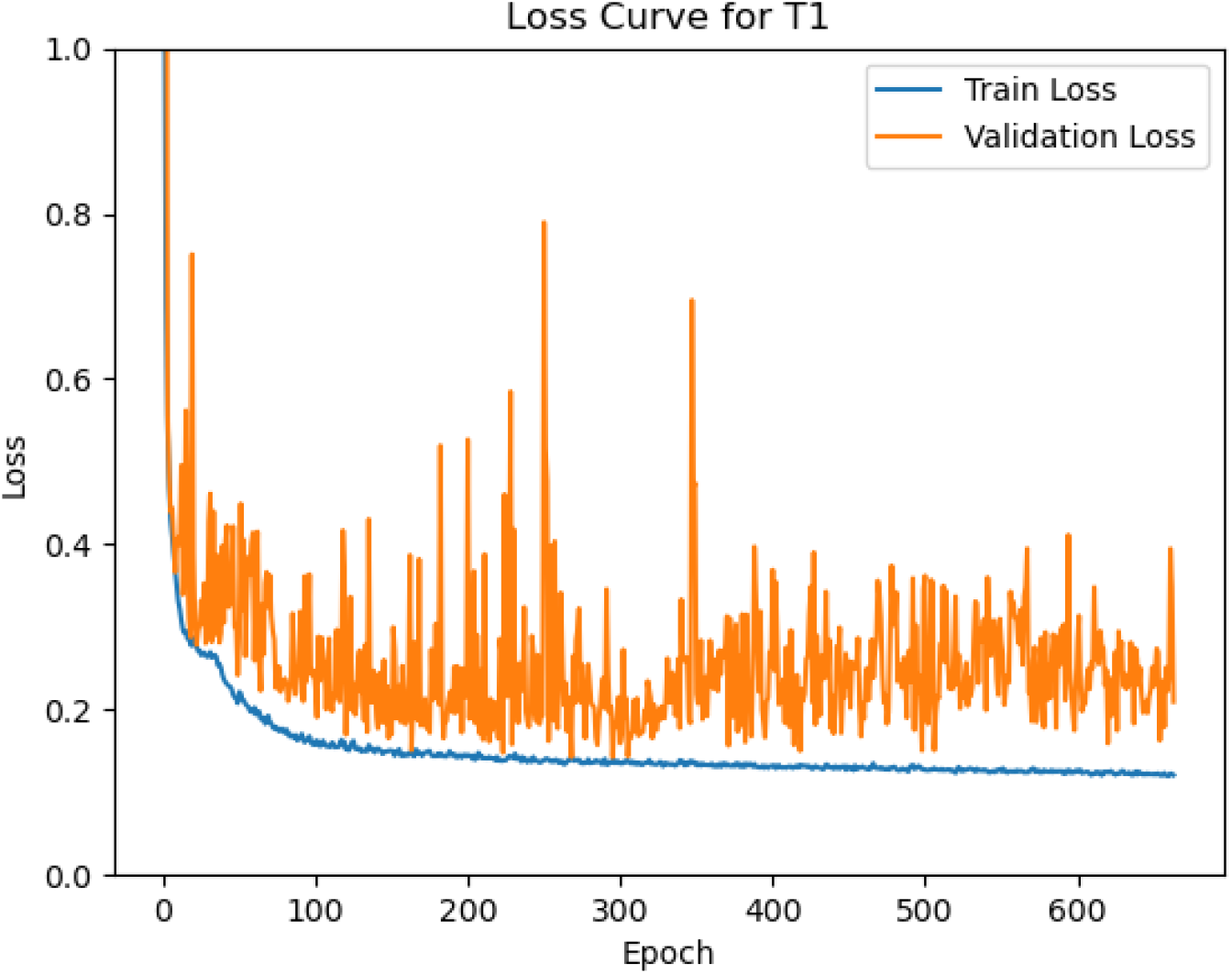
Representative training and tuning loss curves (seed 42). All four models showed similar convergence patterns, reaching early stopping between epochs 663 and 868.

### 3.2 Consensus Performance

The four-model consensus achieved strong performance across evaluation subsets (Table 1). On the held-out test set, the system detected 70 of 73 markers (96% sensitivity [95% CI: 88, 99]) with false positives (95% precision [95% CI: 87, 99]). Across all 34 subjects, 29 (85%) had perfect marker detection.

**Table 1.** Marker-Level Detection Performance of the Four-Model Consensus.

| Subset | TP, FP / Markers | Sensitivity | Precision | Perfect Detection |
| --- | --- | --- | --- | --- |
| Tuning ($n = 9$) | 25, 1 / 26 | 96% (25/26) [80, 100] | 96% (25/26) [80, 100] | 8/9 (89%) |
| Test ($n = 25$) | 70, 4 / 73 | 96% (70/73) [88, 99] | 95% (70/74) [87, 99] | 21/25 (84%) |
| All ($n = 34$) | 95, 5 / 99 | 96% (95/99) [90, 99] | 95% (95/100) [89, 98] | 29/34 (85%) |

Note.—Data are percentages, with numerators and denominators and 95% CIs in brackets. CI = confidence interval, FP = false positive, TP = true positive.

Five subjects had imperfect detection: four had a single nearby-structure substitution (one false negative and one false positive each), and one subject with only two reference standard markers had a spurious third prediction (one false positive, no false negatives).

### 3.3 Model Count Ablation

The consensus approach showed progressive improvement with increasing model count (Fig 4). Individual models showed high variability in sensitivity (74%–96% on the test set), while the four-model consensus reached 96%. The largest improvement occurred from one to two models (+8 percentage points in mean sensitivity). The SD across model combinations decreased from 9% (single models) to less than 1% (three models). Per-subject sensitivity was significantly higher under four-model consensus than under mean single-model inference (mean improvement, +12.2 percentage points; bootstrap 95% CI: 5.8, 18.8; Wilcoxon signed-rank *P* = .002). In 12 of 25 test subjects, consensus produced perfect detection where at least one single model had failed; no subject lost a perfect detection in the opposite direction (sign test *P <* .001).

**Figure 3.**
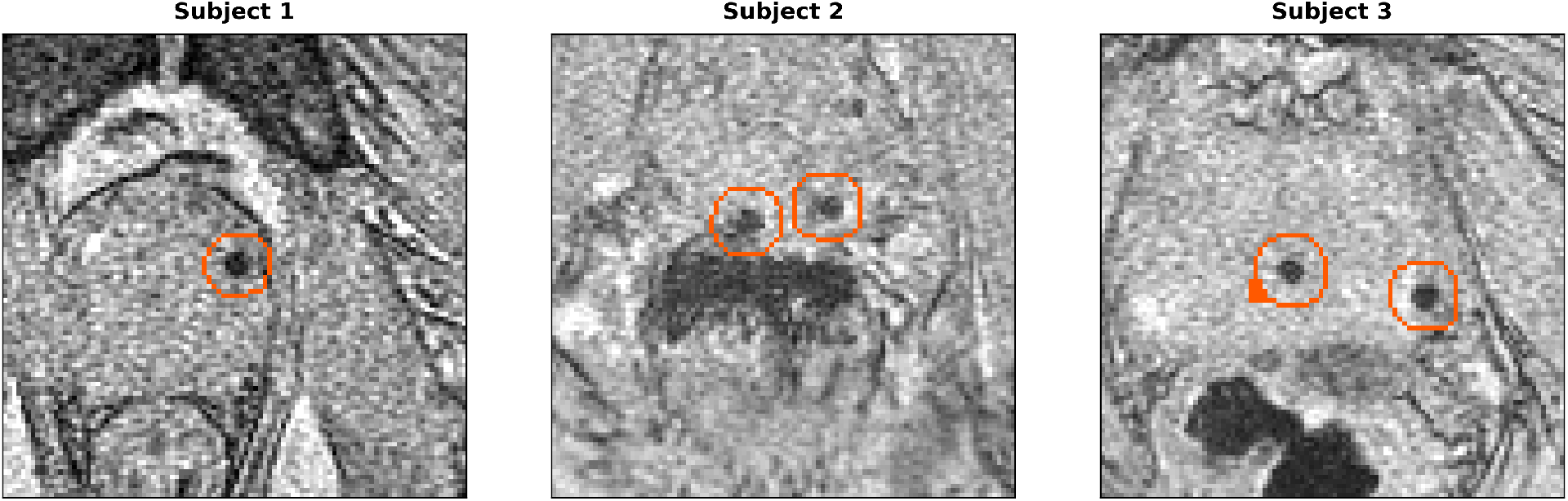
Detected fiducial markers overlaid on T1-weighted axial sections for three test subjects. Orange outlines indicate the boundaries of segmented marker regions. The boundary extraction replicates the on-scanner OpenRecon visualization pipeline (morphological dilation, hole filling, and outer boundary detection).

**Figure 4.**
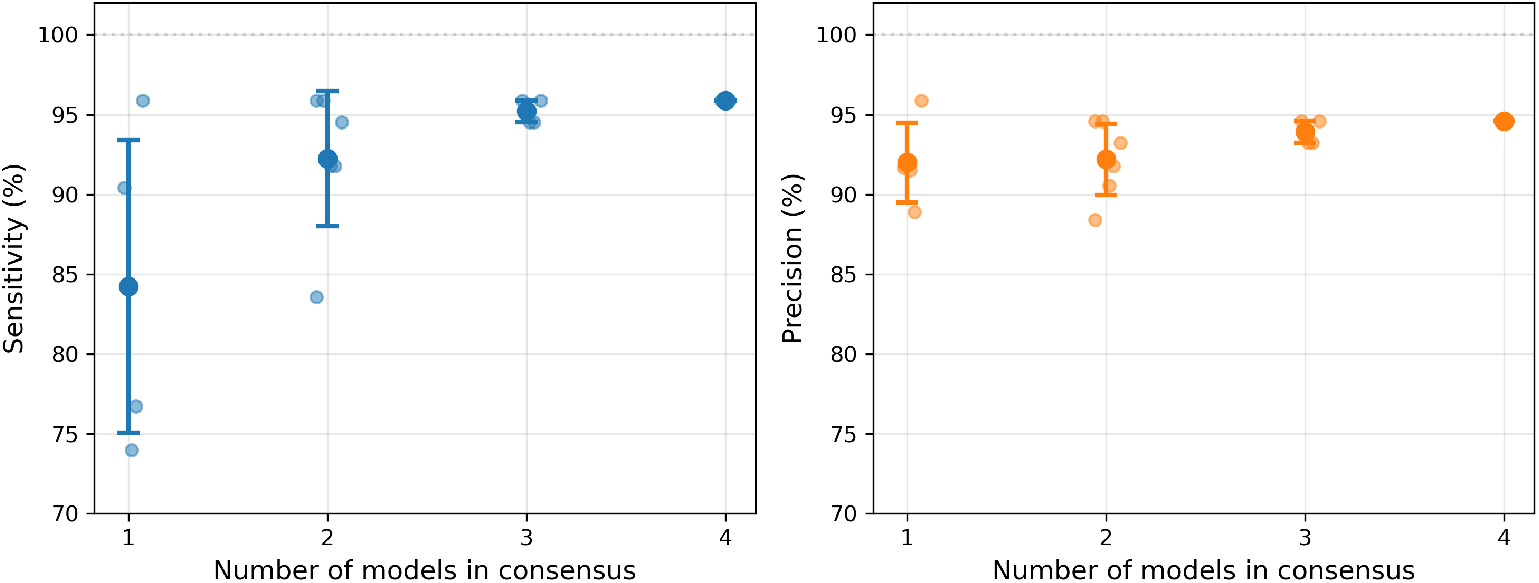
Model count ablation on the test set. Large markers with error bars show mean ± SD across all model combinations; small markers show individual combinations. The consensus progressively improved sensitivity and reduced variability.

Removing the top-3 selection and instead retaining all components above a confidence threshold achieved marginally higher sensitivity (99%; 72 of 73) but substantially reduced precision (80%; 72 of 90) due to numerous false-positive detections.

### 3.4 Failure Analysis

Visualization of failure cases (Fig 5) revealed two prevalent failure modes. In four subjects, the model detected a nearby structure (possibly calcification or tissue heterogeneity) instead of the true FM, resulting in a simultaneous false positive and false negative. In one subject with only two reference standard markers, the model predicted a spurious third marker. These failures involved regions where local appearance closely resembled the marker signature.

**Figure 5.**
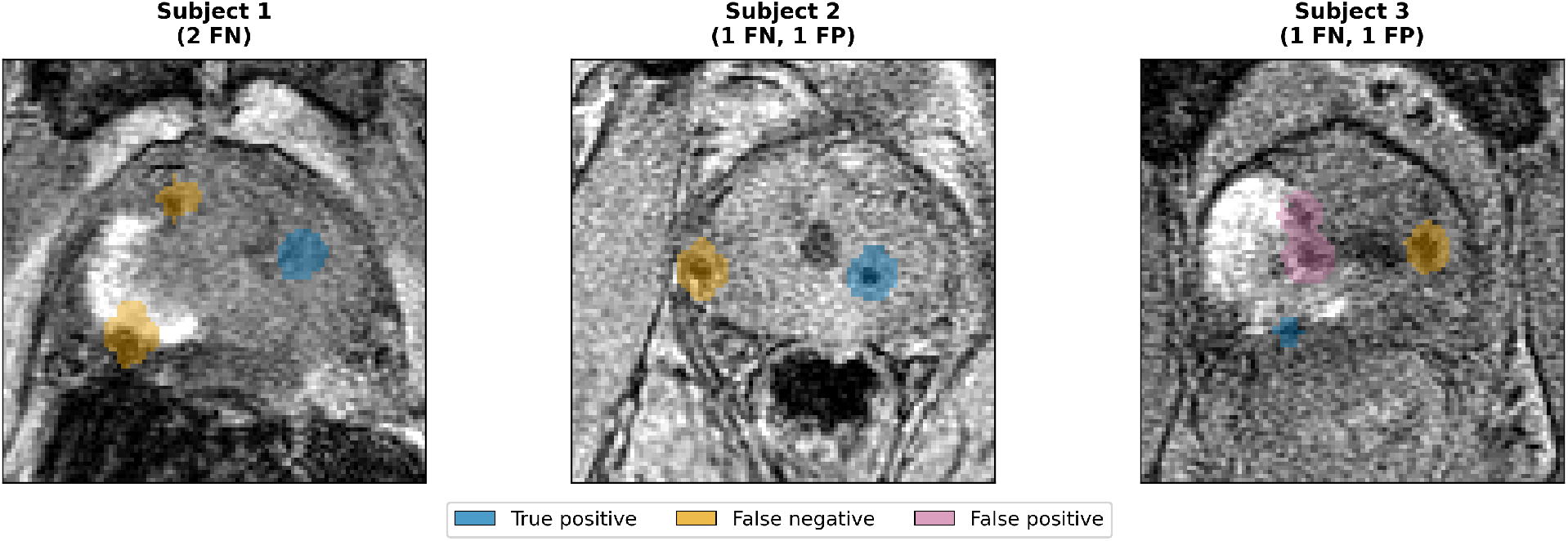
Representative axial slices from three of five subjects with imperfect detection, with colored overlays indicating true positives (blue), false negatives (yellow), and false positives (pink). Each panel shows a single slice; additional markers detected in other slices are not shown. The two omitted subjects had similar failures.

## 4 Discussion

In this study, a four-model consensus deep learning approach using T1-weighted MRI alone achieved 96% sensitivity and 95% precision for automated FM segmentation in 25 held-out test subjects, with 85% of evaluation subjects having perfect marker detection.

The ablation study confirms that consensus significantly improves over single models (*P* = .002), with individual models showing wide performance variability (74%–96% sensitivity) that the consensus stabilizes. Averaging probability maps smooths out individual biases, analogous to test-time augmentation at the model level, while also suppressing spurious activations not consistent across the ensemble. The clinical prior of three markers per patient via top-3 selection was important for precision, as removing this constraint reduced precision to 80%.

Our results are comparable with prior work. Gustafsson et al.^17^ reported 97.4% sensitivity across 39 patients, though with larger FMs (1.2 × 3 mm vs 1 × 3 mm). Other methods using multiparametric MRI achieved 84%–96% true-positive rates.^13–16^ Our approach achieves comparable performance using only routinely acquired T1-weighted images, avoiding additional specialized sequences. Our prior QSM-based approach achieved 80% precision and 90% sensitivity using 26 subjects;^18^ the present consensus approach extends that work with a five-fold larger cohort, routine T1-weighted input, multi-model consensus, and dual deployment pathways.

Beyond segmentation performance, a key contribution is the deployment pipeline bringing the trained models to both the MRI scanner (via OpenRecon) and any modern web browser (via WebAssembly). While many deep learning studies report retrospective metrics, the containerized approach and browser-based application provide practical pathways for clinical integration, with automated continuous integration propagating model updates directly from development to clinical use.

Limitations include that the evaluation set of 34 subjects may not capture the full variability of clinical presentations; uncommon cases such as marker migration or severe calcification were scarce. The temporal split resulted in a slightly older test cohort (mean age, 73 years ± 7 vs 70 years ± 8; *P* = .01, Welch *t* test), though FM appearance on T1-weighted MRI is not expected to vary with age. Reference standard segmentations derived from CT-MRI registration may contain spatial offsets, mitigated by dilation in our evaluation. The fixed top-3 selection assumes exactly three markers; extending the approach to variable counts would improve generalizability. Consensus inference requires four model runs, though processing time remains clinically acceptable (Table 2). Future work should expand the training set with more diverse presentations, particularly edge cases, and pursue prospective multi-site evaluation across scanner manufacturers. Integration with synthetic CT generation could further streamline MR-only radiotherapy workflows.

**Table 2.** Consensus Inference Runtime for a Single Volume (320 × 320 × 40 Voxels) across Deployment Platforms.

| **Platform** | **Time** |
| --- | --- |
| GPU (NVIDIA A100) | $\approx 8$ sec |
| CPU (AMD EPYC 9454, 4 cores) | $\approx 30$ sec |
| Browser (ONNX/WebAssembly, cached) | $\approx 75$ sec |

Note.—GPU and CPU benchmarks were performed on an institutional high-performance computing cluster. Browser time includes bias field correction; first run adds approximately 50 seconds for one-time model download.

## Acknowledgments

We thank the patients involved in this study and the radiation oncology team at Calvary Mater Newcastle, Australia. The authors acknowledge Siemens Healthineers for supporting the project. This research was undertaken with the assistance of resources and services from the Queensland Cyber Infrastructure Foundation (QCIF) and the University of Queensland Research Computing Centre (RCC).

## AI Writing Tools Disclosure

AI-assisted tools (Claude, Anthropic) were used for manuscript editing and formatting. All content was reviewed and verified by the authors.

## References

[1] Rebello RJ, Oing C, Knudsen KE, et al. Prostate cancer. Nat Rev Dis Primers 2021;7(1):1–27. doi: 10.1038/s41572-020-00243-0

[2] Xie Y, Djajaputra D, King CR, Hossain S, Ma L, Xing L. Intrafractional motion of the prostate during hypofractionated radiotherapy. Int J Radiat Oncol Biol Phys 2008;72(1):236–246. doi: 10.1016/j.ijrobp.2008.04.051

[3] Thomas SJ, Ashburner M, Tudor GSJ, et al. Intra-fraction motion of the prostate during treatment with helical tomotherapy. Radiother Oncol 2013;109(3):482–486. doi: 10.1016/j.radonc.2013.09.011

[4] Ng M, Brown E, Williams A, Chao M, Lawrentschuk N, Chee R. Fiducial markers and spacers in prostate radiotherapy: current applications. BJU Int 2014;113(S2):13–20. doi: 10.1111/bju.12624

[5] Schallenkamp JM, Herman MG, Kruse JJ, Pisansky TM. Prostate position relative to pelvic bony anatomy based on intraprostatic gold markers and electronic portal imaging. Int J Radiat Oncol Biol Phys 2005;63(3):800–811. doi: 10.1016/j.ijrobp.2005.02.022

[6] Martin J, Keall P, Siva S, et al. TROG 18.01 phase III randomised clinical trial of the Novel Integration of New prostate radiation schedules with adJuvant Androgen deprivation: NINJA study protocol. BMJ Open 2019;9(8):e030731. doi: 10.1136/bmjopen-2019-030731

[7] Tocco BR, Kishan AU, Ma TM, Kerkmeijer LGW, Tree AC. MR-guided radiotherapy for prostate cancer. Front Oncol 2020;10:616291. doi: 10.3389/fonc.2020.616291

[8] Young T, Dowling J, Rai R, et al. Clinical validation of MR imaging time reduction for substitute/synthetic CT generation for prostate MRI-only treatment planning. Phys Eng Sci Med 2023;46(3):1015–1021. doi: 10.1007/s13246-023-01271-0

[9] Edmund JM, Nyholm T. A review of substitute CT generation for MRI-only radiation therapy. Radiat Oncol 2017;12(1):28. doi: 10.1186/s13014-017-0769-0

[10] Choi JH, Lee D, O’Connor L, et al. Bulk anatomical density based dose calculation for patient-specific quality assurance of MRI-only prostate radiotherapy. Front Oncol 2019;9:997. doi: 10.3389/fonc.2019.00997

[11] Bird D, Nix MG, McCallum H, et al. Multicentre, deep learning, synthetic-CT generation for ano-rectal MR-only radiotherapy treatment planning. Radiother Oncol 2021;156:23–28. doi: 10.1016/j.radonc.2020.11.027

[12] O’Connor LM, Dowling JA, Choi JH, et al. Validation of an MRI-only planning workflow for definitive pelvic radiotherapy. Radiat Oncol 2022;17(1):55. doi: 10.1186/s13014-022-02020-9

[13] Ghose S, Mitra J, Rivest-Hénault D, et al. MRI-alone radiation therapy planning for prostate cancer: automatic fiducial marker detection. Med Phys 2016;43(5):2218–2228. doi: 10.1118/1.4944871

[14] Maspero M, van den Berg CAT, Zijlstra F, et al. Evaluation of an automatic MR-based gold fiducial marker localisation method for MR-only prostate radiotherapy. Phys Med Biol 2017;62(20):7981–8002. doi: 10.1088/1361-6560/aa8975

[15] Dinis Fernandes C, Dinh CV, Steggerda MJ, et al. Prostate fiducial marker detection with the use of multi-parametric magnetic resonance imaging. Phys Imaging Radiat Oncol 2017;1:14–20. doi: 10.1016/j.phro.2017.02.001

[16] Gustafsson C, Korhonen J, Persson E, Gunnlaugsson A, Nyholm T, Olsson LE. Registration free automatic identification of gold fiducial markers in MRI target delineation images for prostate radiotherapy. Med Phys 2017;44(11):5563–5574. doi: 10.1002/mp.12516

[17] Gustafsson CJ, Swärd J, Adalbjornsson SI, Jakobsson A, Olsson LE. Development and evaluation of a deep learning based artificial intelligence for automatic identification of gold fiducial markers in an MRI-only prostate radiotherapy workflow. Phys Med Biol 2020;65(22):225011. doi: 10.1088/1361-6560/ab9f6b

[18] Stewart AW, Goodwin J, Robinson SD, O’Brien K, Jin J, Barth M, Bollmann S. Deep-learning-enabled differentiation between intraprostatic gold fiducial markers and calcification in quantitative susceptibility mapping. bioRxiv 2023. doi: 10.1101/2023.10.26.564293

[19] Tustison NJ, Avants BB, Cook PA, et al. N4ITK: improved N3 bias correction. IEEE Trans Med Imaging 2010;29(6):1310–1320. doi: 10.1109/TMI.2010.2046908

[20] Pérez-García F, Sparks R, Ourselin S. TorchIO: a Python library for efficient loading, preprocessing, augmentation and patch-based sampling of medical images in deep learning. Comput Methods Programs Biomed 2021;208:106236. doi: 10.1016/j.cmpb.2021.106236

[21] Siemens Healthineers. OpenRecon: Open Reconstruction Framework for Magnetic Resonance Imaging. https://www.siemens-healthineers.com/magnetic-resonance-imaging/technologies-and-innovations/openrecon. Accessed May 26, 2026.

[22] NeuroDesk Project. NeuroDesk: A flexible, scalable, and easy to use data analysis environment for reproducible neuroimaging. https://www.neurodesk.org/. Accessed May 26, 2026.

[23] Renton AI, Dao TT, Johnstone T, et al. Neurodesk: an accessible, flexible and portable data analysis environment for reproducible neuroimaging. Nat Methods 2024;21:804–808. doi: 10.1038/s41592-024-02237-2

[24] Inati SJ, Naegele JD, Zwart NR, et al. ISMRMRD: A common data format for MRI raw data. J Magn Reson Imaging 2017;45(6):1767–1774. doi: 10.1002/jmri.25428

[25] ONNX Runtime developers. ONNX Runtime. https://onnxruntime.ai/. Accessed May 26, 2026.

[26] Eckstein K, Dymerska B, Bachrata B, et al. MriResearchTools: A Julia package for MRI data processing. Magn Reson Med 2024;92(4):1498–1508. doi: 10.1002/mrm.30170

[27] Li X, Morgan PS, Ashburner J, Smith J, Rorden C. The first step for neuroimaging data analysis: DICOM to NIfTI conversion. J Neurosci Methods 2016;264:47–56. doi: 10.1016/j.jneumeth.2016.03.001

[28] Rorden C, Webster M, Drake T, Hanayik T. NiiVue: a WebGL2 medical image viewer. Front Neuroinform 2024;18:1344312. doi: 10.3389/fninf.2024.1344312

[29] Stewart AW, Bollmann S. dicompare: Validation of medical imaging sequences to facilitate collaboration across global sites. https://github.com/astewartau/dicompare. Accessed May 26, 2026.

